# The Antimicrobial Activity of *Pechuel-Loeschea leubnitziae* Leaf Extract and its Effect on The Expression Level of Methicillin Resistant *Staphylococcus Aureus* and *Candida albicans* Virulence-Associated Genes

**DOI:** 10.1101/2022.01.11.475881

**Authors:** Mariah Ndilimeke Muhongo, Mourine Kangogo, Christine Bii

## Abstract

The complete halt in the synthesis of new effective antimicrobial compounds is a global concern. Pathogenic microorganisms’ virulence mechanisms seem to have a significant impact on their pathogenesis. The purpose of this study was to examine the antimicrobial activity of the ethanol and methanol fractions of *Pechuel-Loeschea leubnitziae* leaf extract, as well as its effect on the expression level of virulence-associated genes.

The extract’s fractions were evaluated for antimicrobial activity against *Escherichia coli* ATCC 25922, Pseudomonas aeruginosa ATCC 27853, *Klebsiella pneumoniae* (clinical), Methicillin-resistant *Staphylococcus aureus*, and *Candida albicans* ATCC 90029. The test organism’s antibiogram pattern was determined. The extracts’ attenuation effect on the target genes of the susceptible organisms was investigated employing relative quantification using RT-qPCR. The test organism’s antibiogram pattern revealed that it was drug-resistant, intermediate, and sensitive. The extracts tested positive for antimicrobial activity against Methicillin-resistant *Staphylococcus aureus* and *Candida albicans* ATCC 90029, with zones of inhibition varying from 20.33 to 29 mm. The lowest recorded MIC value was 4.688 mg/ml, while the highest was 37.5 mg/ml. In contrast to the methanol extract, the ethanol extract had a cidal action at a lower dose. The ethanol extract’s Sub-MIC (18.25 mg/ml) merely reduced the expression of the hly gene in MRSA. The MRSA virulence genes were not suppressed by the sub-MIC of methanol extract (18.25 mg/ml). Notably, the expression of als1, pbl1, and sap1 in *Candida albicans* ATCC 90029 was significantly attenuated when exposed to sub-MICs of ethanol extract (2,344 mg/ml) and methanol extract (9.375 mg/ml). Per the findings of this research, the leaves of *P. leubnitziae* could be a source of an effective antimicrobial agent in the therapy of MRSA/*Candida*-related disorders.

## INTRODUCTION

Antimicrobial resistance is one of the greatest threats to human health globally, causing an estimated 700 000 deaths annually [1]. It is predicted that by 2050, there will be approximately 10 million deaths [2]. Antimicrobial resistance is worsening in many African and Asian countries as a result of high levels of counterfeit and substandard antimicrobials in the pharmaceutical market [3]. Antimicrobial-resistant bacterial infections kill nearly 50 000 people each year in Europe and the United States alone, with hundreds of thousands more dying in other parts of the world [4].

According to a WHO circular, the antimicrobial resistance estimates in *E. coli* and *S. aureus* are of the most significant concern. They are listed as the most common cause of hospital-associated infections. Previous studies indicated at least 50 % resistance to fluoroquinolones and third-generation cephalosporins among *E. coli* isolates [1]. According to [1], the circular cited *Escherichia coli, Pseudomonas aeruginosa, Klebsiella pneumoniae*, Methicillin-resistant *Staphylococcus aureus* and, *Candida albicans* as among the common pathogens associated with multi-drug resistance.

Since many pathogens are becoming drug resistant, there is indeed a keen interest in natural remedies such as medicinal plants. Medicinal plants have been exploited for millennia by diverse societies around the world to heal various ailments [5, 6]. The consumption of herbal remedies, particularly in impoverished countries such as Namibia, greatly contributes to basic health care since they are readily available and inexpensive. According to numerous surveys done by WHO, around 80% of the world’s population relies on traditional medicine. There are approximately 350,659 medicinal plants in the world, yet only 2% have been explored and reported based on their therapeutic biological properties [2]. Most medicinal plants in Namibia are amply documented in the botanical and are prominent due to their rich heritage of being used. Despite the reality that there is sufficient knowledge on their indigenous application, there is still a massive disparity in the scientific evidence of their bioactive compounds content and biological efficacy [7, 8].

*Pechuel-Loeschea leubnitziae* is a species of the *Asteraceae* family. It is a harsh shrub that is popularly known as a bitter bush in English, however its local name in Oshiwambo is Oshizimba [9]. *Pechuel-Loeschea leubnitziae* is a plant species which may be spotted across Namibia. This has traditionally been used to alleviate gastrointestinal problems, venereal diseases, and pharyngitis [10]. To scientifically authenticate a medicinal plant’s traditional use, its biological potency must be determined. Thus, evaluating the antimicrobial activity of the *Pechuel-Loeschea leubnitziae* leaf extract and its effects on the expression of major microbial virulence genes is relevant [11].

Per this background, the research was planned to screen fractions of *Pechuel-Loeschea leubnitziae* leaf extract for antimicrobial activity against selected pathogens of public health importance, as well as to assess its effect on virulence-associated genes of various virulence factors. The agar well diffusion procedure was used for antimicrobial assay, and the relative quantification approach was used for the gene expression analysis. The antimicrobial activity of leaf extracts was contrasted to the effects of conventional antibiotics, and the CT values of the target genes were normalized with endogenous genes (16s rRNA for MRSA) and (18s rRNA for Candida albicans ATCC 90029), and the expression of the target genes in treated samples was compared to untreated samples.

## MATERIALS AND METHODS

### Sample Preparation and Extraction

The leaves were properly rinsed with water and air-dried. To obtain a fine powder for the extraction process, the dry sample was pulverized with a household blender. In this study, the cold maceration technique was combined with the serial exhaustive extraction approach. The extract was prepared by soaking 360g in 1800 ml of Ethanol (1:5 ratio) on a shaker for 72 hours. The experiment was repeated with Methanol on the residual. The resultant samples were filtered with two layers of muslin cloth before being gravitational filtered through Whatman number 1 filter papers. The filtrates were concentrated using a rotatory evaporator at temperatures ranging between 40 to 60 °C [12]. The extracts were then stored at 4 °C for subsequent analysis.

### Collection and Maintenance of Test Organism

The isolates were obtained from Kenya Medical Research Institute (KEMRI). *Escherichia coli* ATCC 25922, *Pseudomonas aeruginosa* ATCC 27853, *Klebsiella pneumoniae* (clinical), Methicillin-resistant *Staphylococcus aureus*, and *Candida albicans* ATCC 90029 were the microbial isolates used. The colonies of the test microorganisms were sub-cultured on nutrient agar.

### Antimicrobial Susceptibility Test

The agar well diffusion susceptibility method was used to evaluate the ethanol and methanol extracts for antimicrobial activity, according to the Clinical and Laboratory Standards Institute [13]. The isolates of the test organisms were sub-cultured on nutrient agar plates and incubated for 18 hours at 37 °C. The turbidity of the cultures was adjusted to 0.5 Macfarland standard (108 Cfu/ml) using normal saline. A sterile cotton swab was dipped in the standardized suspension and placed on the surface of Mueller Hinton Agar plates and allowed to diffuse for five minutes. A sterile cork borer with a 6 mm diameter was used to make the agar wells. Exactly 50 μl of varying concentrations of each solvent extract was introduced into the wells. The Antibiotic discs used for *Escherichia coli* ATCC 25922 and *Klebsiella pneumoniae* (clinical) are amikacin (30 μg amoxicillin-clavulanate (10 μg), ampicillin (10 μg), cefepime (30 μg), and gentamicin (10 μg). For *Pseudomonas aeruginosa* 27853 are amikacin (30 μg), cefepime (30 μg), and gentamicin (10 μg). For Methicillin-Resistant *Staphylococcus aureus* are cefoxitin (30 μg), oxacillin (5 μg), erythromycin (15 μg) and gentamicin (10 μg) while for Candida albicans ATCC 90029 are fluconazole (100IU) and Nystatin (100IU). As a control, 5% DMSO was employed. The plates were allowed to diffuse for about 15 minutes at room temperature before being incubated at 37 °C for 18 hours. Following incubation, the plates were examined for the formation of a clear zone around the wells, which correlates with the antimicrobial activity of the tested solvent extracts. The diameter of the zones of inhibition was measured in mm using a ruler in triplicate.

### Minimum Inhibitory Concentration (MIC) Analysis

The ethanol and methanol extracts’ Minimum Inhibitory Concentration (MIC) on susceptible organisms was assessed using broth microdilution according to [13] approved guidelines. The 100 μl of initial concentration (600 mg/ml) of each solvent extract was diluted with 100 μl of Muller Hinton broth in a 96-well round-bottom microtiter plate using double-fold serial dilution to obtain ten different concentrations: (600, 300, 150, 75, 37.5, 18.75, 9.375, 4.688, 2.344, and 1.172 mg/ml). A standardized (0.5 McFarland) inoculum of 10 μl was added and stirred. MHB plus inoculum, MHB plus extract, and MHB plus 5% DMSO solution were used as negative controls. As a positive control, conventional drugs were used: ampicillin (50 g/ml) for MRSA and nystatin (100IU) for Candida albicans ATCC 90029. The plate was wrapped with a sterile parafilm and incubated for 24 hours at 37 °C. Following incubation, 20 μl of 0.05% resazurin dye was added to the wells and incubated for another 2 hours to examine colour patterns. Wells with microbial growth will turn pink, while wells without microbial growth will remain blue. The MIC was defined as the lowest extract concentration that totally inhibited microbial growth.

### Minimum Bactericidal Concentration (MBC) Analysis

Sub-culturing the wells of Methicillin resistant *Staphylococcus aureus* that showed no colour change upon post-incubation of the microtiter plate yielded the MBC of each solvent extract. This was done by dipping a sterile inoculation loop into each well and infecting on Mueller Hinton agar plates. The plates were maintained for 18 hours at 37 °C. The MBC was determined to be the lowest concentration that showed no visible growth after incubation.

### Minimum Fungicidal Concentration (MFC) Analysis

Sub-culturing the wells of *Candida albicans* ATCC 90029 that showed no colour change upon post-incubation of the microtiter plate yielded the MFC of each solvent extract. This was done by dipping a sterile inoculation loop into each well and infecting on Mueller Hinton agar plates. The plates were maintained for 18 hours at 37 °C. The MFC was determined to be the lowest concentration that showed no visible growth after incubation.

### RNA Extraction and cDNA Preparation

The samples were prepared by culturing 100 μl of 0.5 Macfarland standardized (108 Cfu/ml) of MRSA and *C. albicans* ATCC 90029 in 4 ml of Tryptic soy broth and incubated with or without 100 μl of sub-MICs concentration of ethanol (18.25 and 2.344 mg/ml) and methanol (18.25 and 9.375 mg/ml) extracts for 16 hours at 37 °C on a shaker with 160 rpm. The cells were harvested by centrifugation of 1 ml of the suspension (13,000 × g, 5 minutes, 4 °C), and discarded the supernatant. The cells were rinsed with chilled PBS three times at (350 × g, 5 minutes, 4 °C). The cells were resuspended in 200 μl of RNAse free water and sonicated five times for 10 seconds. Total RNA was extracted using PureLinkTM RNA Mini kit (Invitrogen by Thermo Fisher Scientific) based on the manufacturer’s protocol. Each sample was treated with DNase I according to the manufacturer’s instructions to remove any possible DNA contamination. The concentration and purity of each RNA sample were determined using a NanoDrop spectrophotometer (PCRmax Lambda) at 260/280 and 260/230 nm. The RNA integrity was analyzed by gel electrophoresis on a 1 % agarose gel. The 0.1 μg of RNA template was reverse-transcribed into complementary DNA (cDNA) using a FIREScript RT cDNA synthesis kit according to the manufacturer’s instructions. Samples were stored at -20 °C until further analysis.

### Gene expression analysis by RT-qPCR

The primers were designed using the PRIMER3 tool version (0.4.0). The amplification specificity was validated with an *in-silico* PCR amplification and Sequence Manipulation Suite (SMS). The primers were synthesized by Macrogen, Europe. The details of the primers are listed in **Table 1**. For gene expression analysis, real-time-quantitative polymerase chain reaction (RT-qPCR) was performed using a q-Tower3 84 (Analytik Jena) and Luna® Universal qPCR Master Mix (M300L). Each 20 μl PCR reaction contained 10 μl of Luna universal qPCR Master 1X mix, 0.25 μM of each forward and reverse primers and about 2 μl of 100x diluted cDNA. The PCR was set to initial denaturation at 95 ºC for 60 seconds, 40 cycles of denaturation at 95 ºC for 15 s, and annealing at 55 ºC for 30 s. At the end of the PCR, a melting curve program was set from 60 ºC to 95 ºC with a 0.5 ºC increase every 15 s. All samples were analyzed in triplicate. The housekeeping genes (16s rRNA and 18s rRNA) were used as internal controls to normalize the expression level of the samples.

**Table 1.**
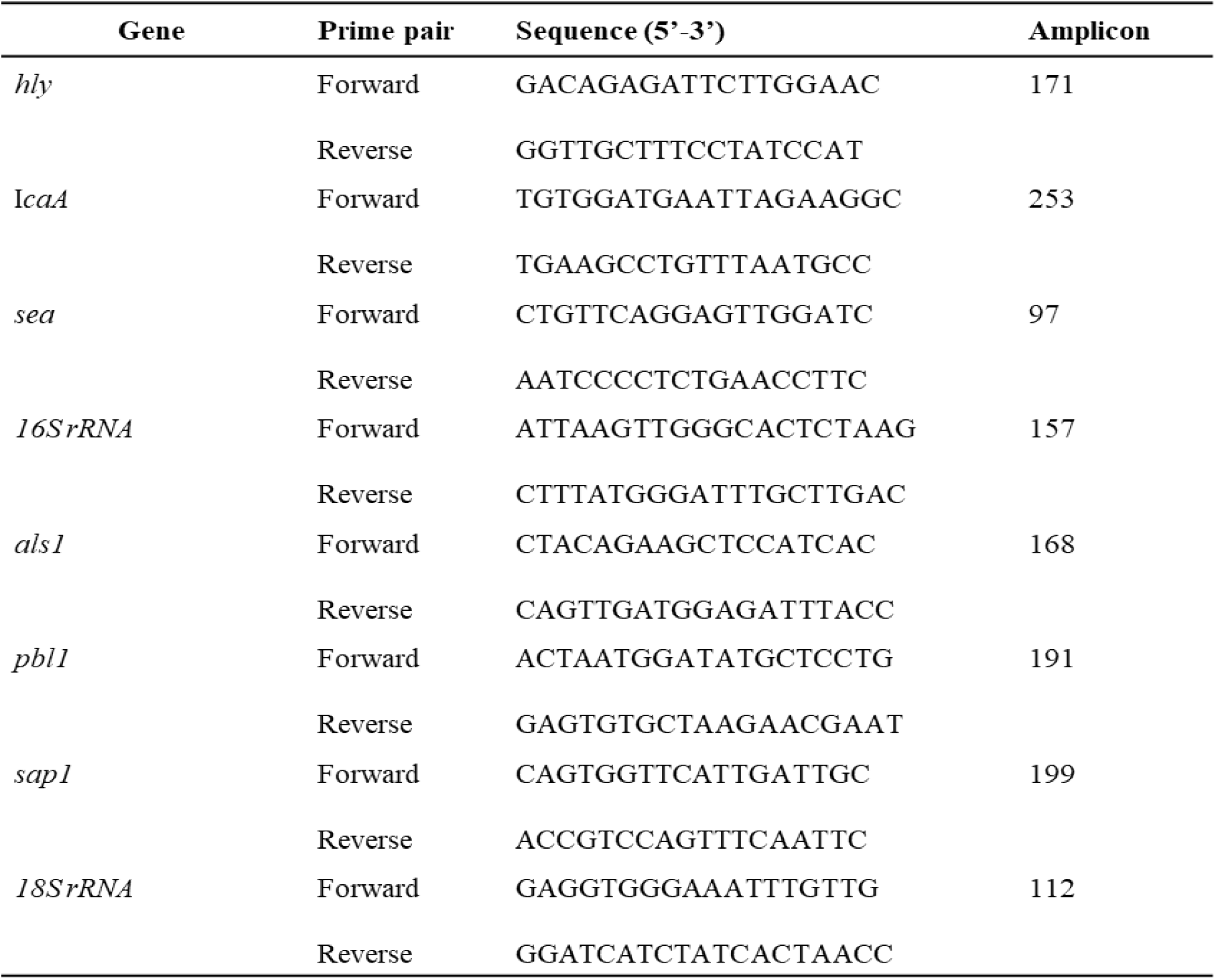
Primers used for expression of virulence genes and reference genes (16s rRNA and 18s rRNA).

### Data analysis

All data were captured in Microsoft® Excel spreadsheets. Data from the experiment were analyzed using SPSS (20.1 version). All results were done in triplicate and presented as the mean ± standard deviation (S.D.). The MIC, MBC, and MFC were analyzed using descriptive statistics. The expression levels of the target genes were measured relative to the control sample and normalized to the expression of the endogenous reference gene (16s/ 18s ribosomal RNA gene) between samples in parallel. The relative expression level or *n*-fold change of target genes between treated and non-treated were calculated using the 2-ΔΔct method according to [14]. The normalized expression levels of the genes in treated and untreated samples were compared using an unpaired two-tailed *t*-test at a significant level of *p* < 0.05. All values of n-fold differential expression were plotted in a graph. Graphs were prepared using GraphPad Prism version 8.3.4 and SigmaPLot version 14.0.

## RESULTS

### Antibiotic Susceptibility Test

The AST zones of inhibition were evaluated in accordance with the Clinical Laboratory Standard Institute [13]. Amikacin, cefepime, and gentamicin were all effective against *E. coli* ATCC 25922, *P. aeruginosa* ATCC 27853, and *K. pneumoniae* (clinical). The *E. coli* ATCC 25922 strain was sensitive to amoxicillin-clavulanate (19.33 mm) but partially resistant to ampicillin (16.33 mm). Additionally, *K. pneumoniae* (clinical) was amoxicillin-clavulanate sensitive (22.33 mm) but ampicillin resistant. The Methicillin-Resistant *Staphylococcus aureus* tested positive for gentamicin (25.67 mm) but was resistant to cefoxitin, oxacillin, and erythromycin. *Candida albicans* ATCC 90029 was completely resistant to nystatin and fluconazole (**Figure 1**).

**Figure 1.**
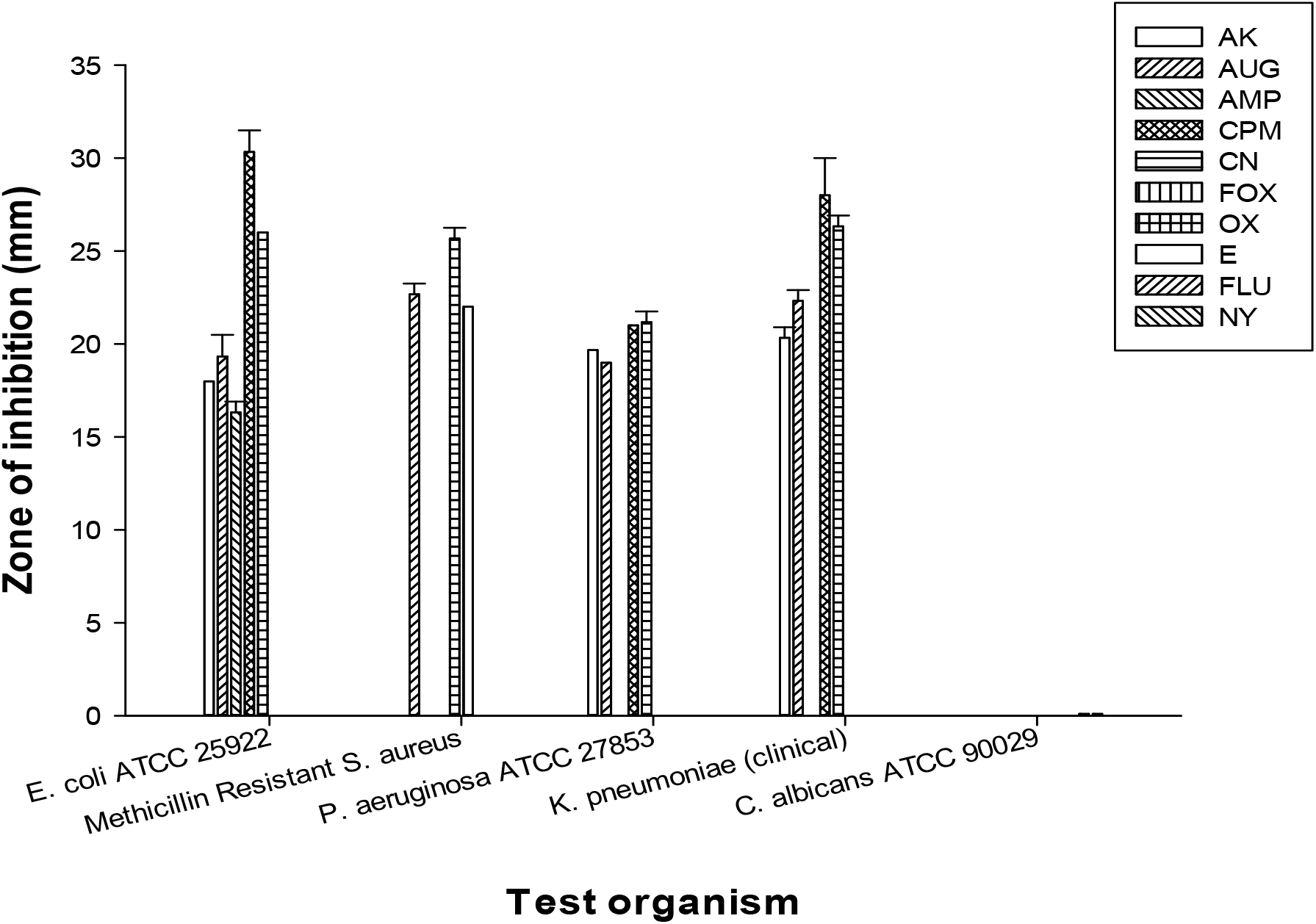
Antibiogram profile of the test organisms. *Key: AK - Amikacin, AUG - Amoxicillin-clavulanate, AMP - Ampicillin, CPM - Cefepime, CN - Gentamicin, FOX - Cefoxitin, OX - Oxacillin, E - Erythromycin, FLU - Fluconazole, NY - Nystatin. Key: Results are expressed as means ± standard deviation of triplicates*.

### Antimicrobial Susceptibility of *Pechuel-Loeschea leubnitziae* Leaf Extract

The zones of inhibition data were reported. The leaf extract of *Pechuel-Loeschea leubnitziae* was tested for antimicrobial activity against multi-drug resistant pathogens at concentrations of 200, 400, and 600 mg/ml. The ethanol and methanol extracts exhibited no inhibitory effect on *Escherichia coli* ATCC 25922, *Pseudomonas aeruginosa* ATCC 27853, and *Klebsiella pneumoniae* (clinical), according to the results of the zones of inhibition. However, the extracts inhibited *C. albicans* ATCC 90029 at all concentrations, with zones of inhibition varying between 22 to 29.33 mm. Additionally, at the highest concentration (600 mg/ml), the extracts inhibited MRSA growth (**Figures 2 and 3**).

**Figure 2.**
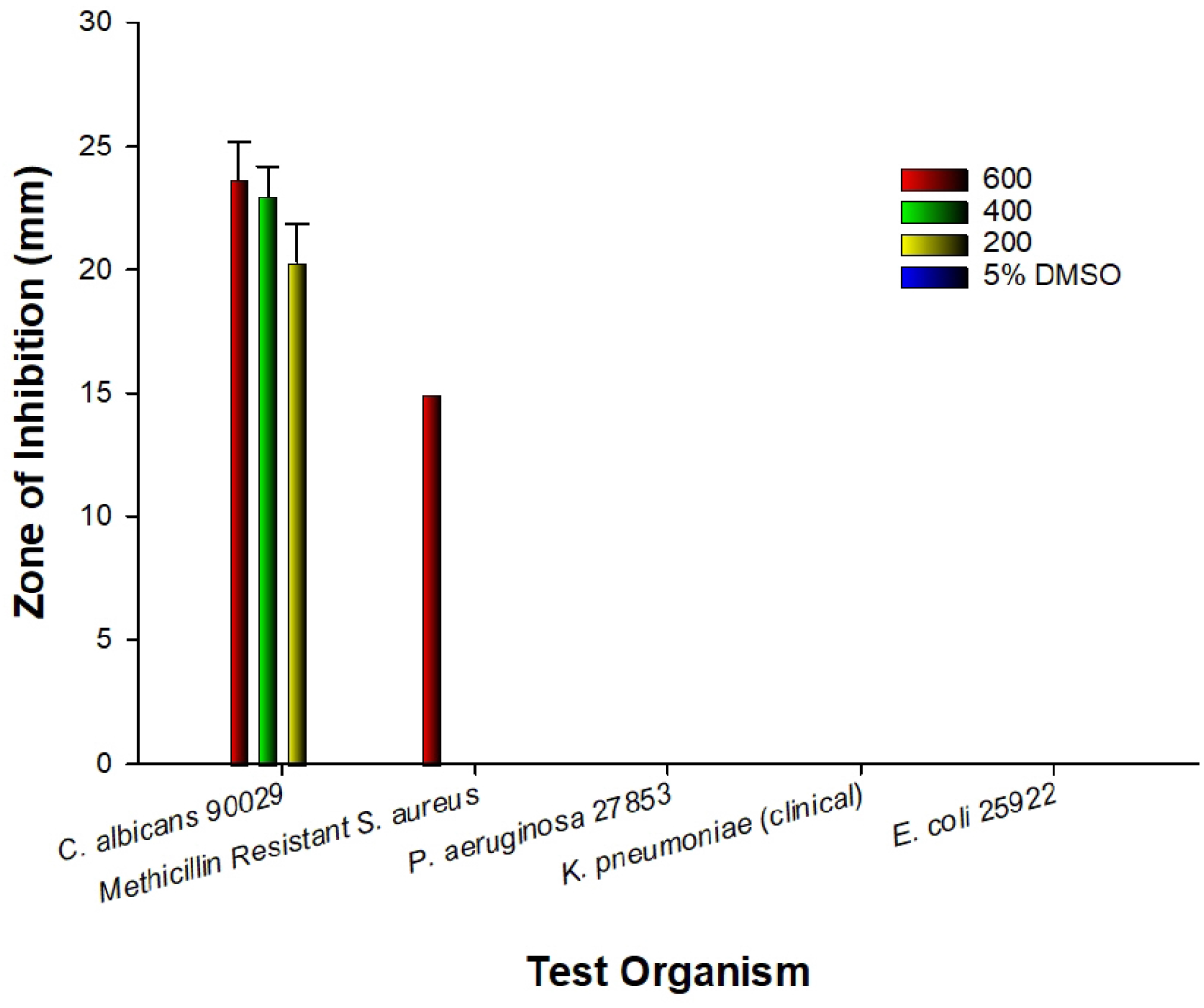
Susceptibility of the test organisms to ethanol extract of *P. leubnitziae* *Key: Results are expressed as means ± standard deviation of triplicates*.

**Figure 3.**
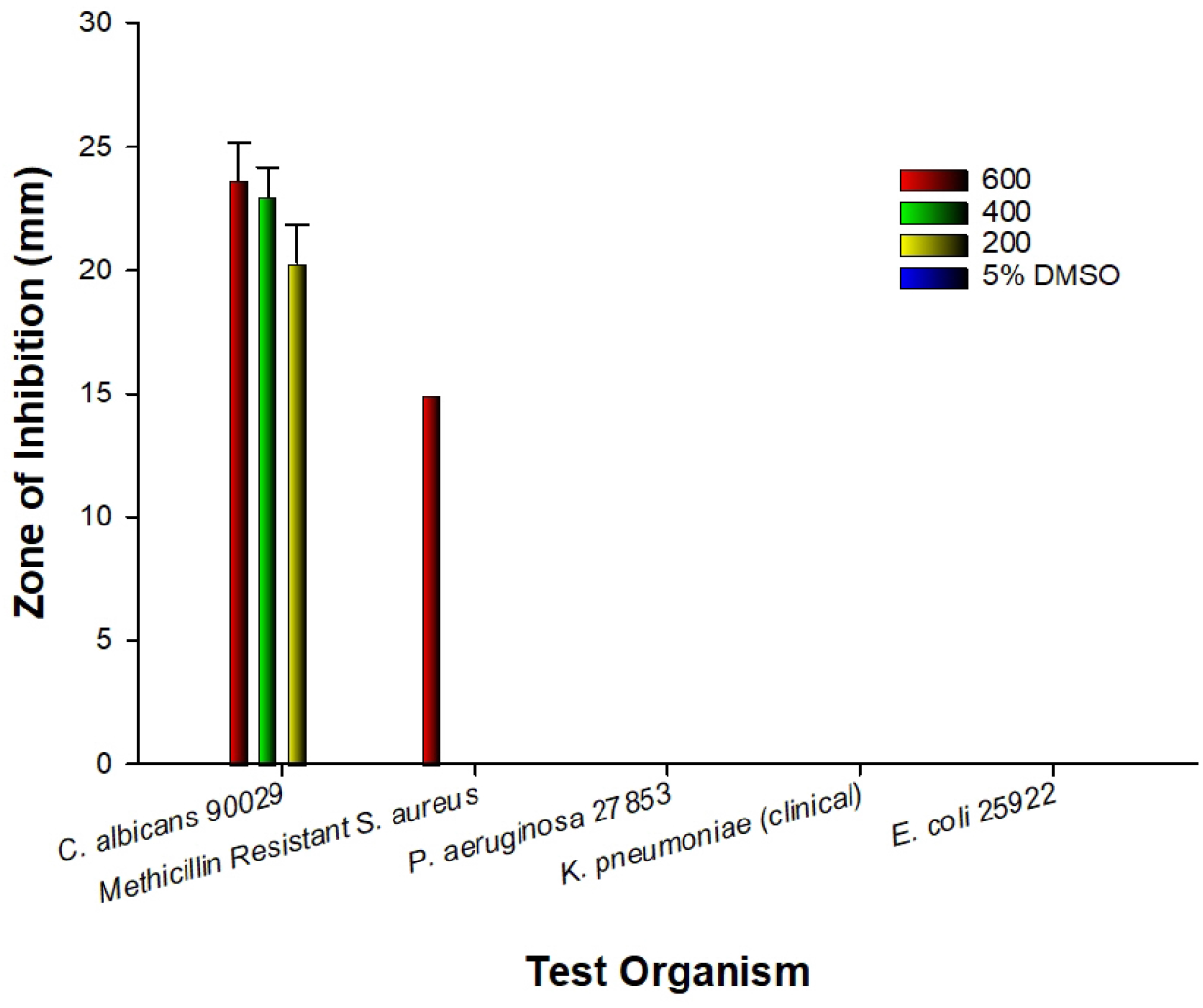
Susceptibility of the test organisms to methanol extract of *P. leubnitziae* *Key: Results are expressed as means ± standard deviation of triplicates*.

### Minimum Inhibitory and Bactericidal/ Fungicidal Concentrations of the Ethanol and Methanol Extracts of *Pechuel-Loeschea leubnitziae* Leaf

By observing the wells without microbial growth (wells that remained blue after the addition of resazurin), the lowest inhibitory concentration of the ethanol and methanol extracts was determined. The lowest inhibitory concentration of the ethanol extract was found to be 37.5 mg/ml for Methicillin-Resistant *Staphylococcus aureus* and 4.688 mg/ml for *Candida albicans* ATCC 90029 (**Table 2**). The MIC of the methanol extract was 37.55 mg/ml for Methicillin-Resistant *Staphylococcus aureus* and 18.75 mg/ml for *Candida albicans* ATCC 90029 (**Table 3**). The ethanol extract exhibited bactericidal and fungicidal action against Methicillin-Resistant *Staphylococcus aureus* and *Candida albicans* ATCC 90029 at concentrations of 37.55 mg/ml and 18.75 mg/ml, respectively. The bactericidal and fungicidal actions were observed in both test organisms at a concentration of 37.5 mg/ml of methanol extract.

**Table 2.**
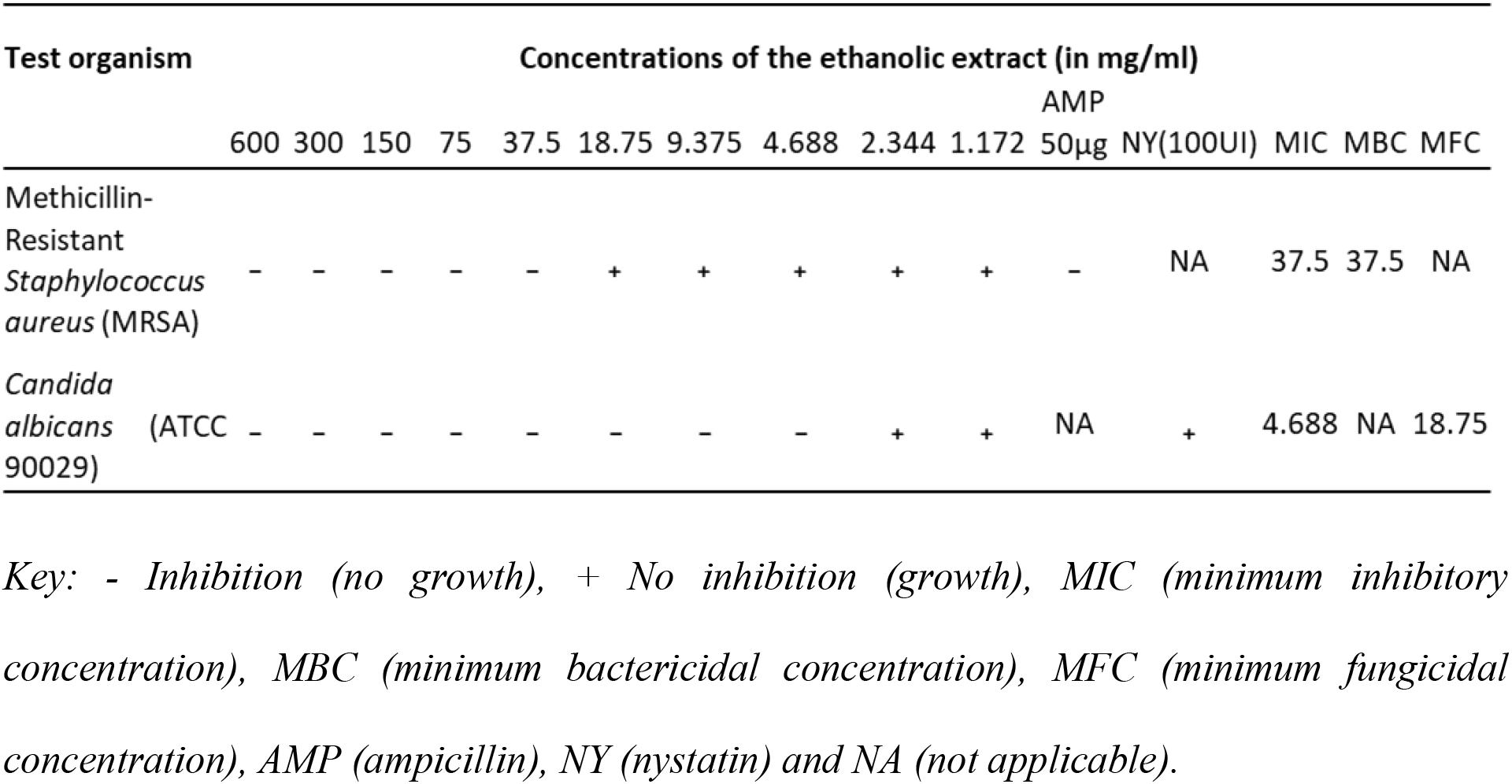
Minimum inhibitory and bactericidal/fungicidal concentrations of the ethanol extract.

**Table 3.**
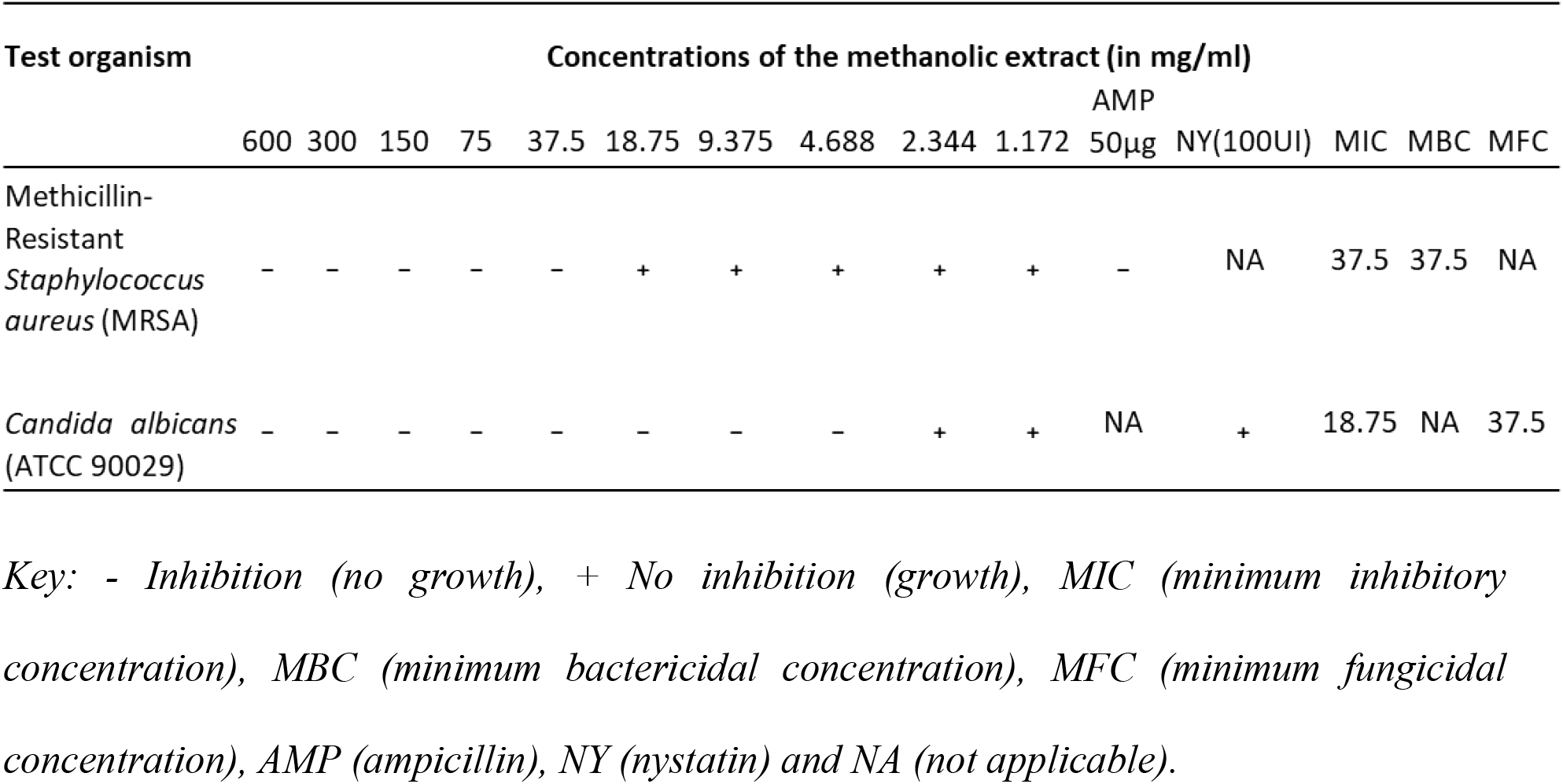
Minimum inhibitory and bactericidal/fungicidal concentrations of the methanol extract.

### Agarose Gel Electrophoresis of Total RNA

The gel electrophoresis (agarose 1%) stained with Gel red dye was used to evaluate the integrity of the RNA samples. A UV gel documentation system was used to visualize and generate the gel images. The total RNA was isolated from treated and untreated MRSA samples (**Figure 4**) as well as C. albicans ATCC 90029 samples (**Figure 5**).

**Figure 4.**
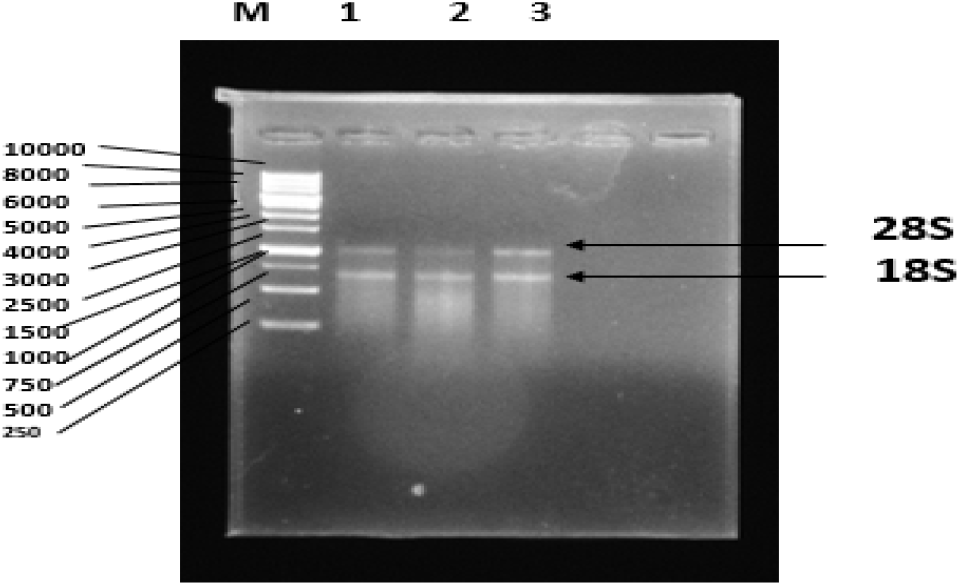
Gel electrophoresis of MRSA RNA samples. *Key: M-marker (1kb). 1-Ethanol-treated sample, 2-Methanol-treated sample, and 3-Untreated sample*.

**Figure 5.**
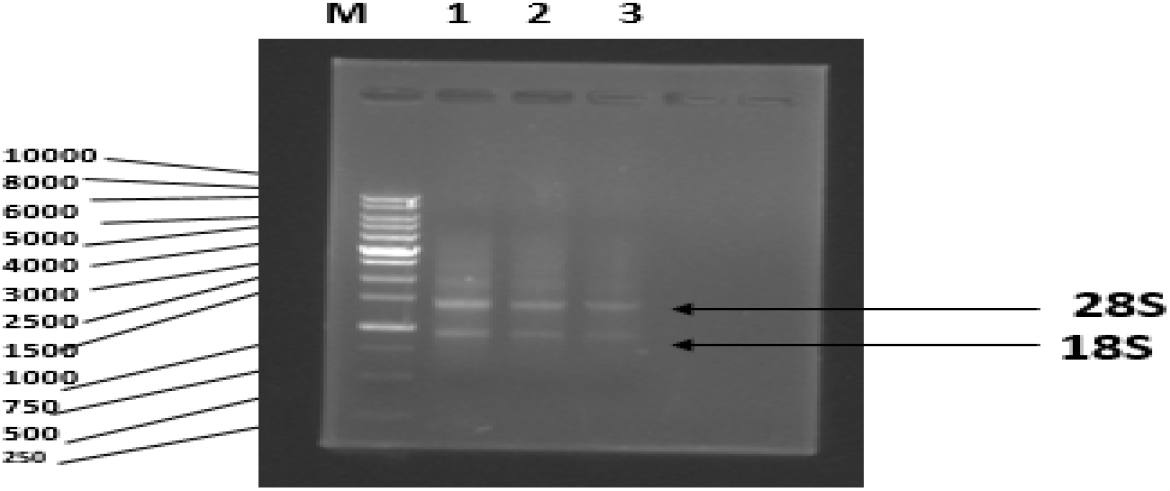
Gel electrophoresis of *C. albicans* ATCC 90029 RNA samples. *Key: M-marker (1kb). 1-Untreated sample, 2-Ethanol-treated sample, and 3-Methanol-treated sample*.

### Validation of the Housekeeping Genes (16S rRNA and 18S rRNA)

The use of housekeeping genes in the calibration of target gene expression in both control and treated samples was validated by calculating the statistically significant difference between samples using ANOVA. The results of the 16S rRNA experiment were formulated. There was no significant difference in the mean CT values (*p*=0.5354) between treated and untreated gene expression values (**Table 4**). The mean CT values ± S.D. was (13.87±0.258, 13.53±0.331, and 13.72±0.448) for the control, EtOH-E treatment, and MeOH-E treatment, respectively (**Figure 6**). The 18S rRNA result was reported (**Table 5**). There was no significant difference in the mean CT values (p=0.0813). The mean CT values ± S.D. of the control, EtOH-E treatment, and MeOH-E treatment was (16.78±0.180, 16.61±0.301, and 17.11±0.056) respectively (Figure 7).

**Table 4.** The 16S rRNA gene’s one-way ANOVA results for the control, EtOH-E treatment, and MeOH-E treatment.

|  | ANOVA | SS | DF | MS | F (DFn, DFd) | P value |
| --- | --- | --- | --- | --- | --- | --- |
|  | Treatment (between groups) | 0.1742 | 2 | 0.0871 | F (2, 6) = 0.6946 | P=0.5354 |
|  | Within group | 0.7523 | 6 | 0.1254 |  |  |
|  | Total | 0.9265 | 8 |  |  |  |

| Tukey's multiple comparisons test | Mean Diff. | 95% CI of diff. | P Value | Summary |
| --- | --- | --- | --- | --- |
| Control vs. EtOH-E treatment | 0.340 | -0.5471 to 1.227 | 0.5079 | ns |
| Control vs. MeOH-E treatment | 0.150 | -0.7371 to 1.037 | 0.8652 | ns |
| EtOH-E treatment vs. MeOH-E treatment | -0.190 | -1.077 to 0.6971 | 0.7954 | ns |

|  | Control | EtOH-E | MeOH-E |
| --- | --- | --- | --- |
| Mean ct | 13.87 | 13.53 | 13.72 |
| Std. Dev | 0.258 | 0.331 | 0.448 |
| SEM | 0.149 | 0.191 | 0.259 |

Key: ANOVA- Analysis of Variance, SS- the sum of the square, DF- degree of freedom, SEM- standard error of the mean, EtOH-E (ethanol extract), and MeOH-E (methanol extract).

**Table 5.** The 18S rRNA gene’s one-way ANOVA results for the control, EtOH-E treatment, and MeOH-E treatment

|  | ANOVA | SS | DF | MS | F (DFn, DFd) | P value |
| --- | --- | --- | --- | --- | --- | --- |
|  | Treatment (between groups) | 0.3241 | 2 | 0.1621 | F (2, 4) = 5.015 | P=0.0813 |
|  | Within group | 0.1293 | 4 | 0.03232 |  |  |
|  | Total | 0.4534 | 6 |  |  |  |
| Tukey's multiple comparisons test | Mean Diff. | 95% CI of diff. | P Value | Summary |  |  |
| Control vs. EtOH-E treatment | 0.170 | -0.4707 to 0.8107 | 0.6442 | ns |  |  |
| Control vs. MeOH-E treatment | -0.330 | -0.9149 to 0.2549 | 0.225 | ns |  |  |
| EtOH-E treatment vs. MeOH-E treatment | -0.500 | -1.085 to 0.08488 | 0.0798 | ns |  |  |
|  |  | Control | EtOH-E | MeOH-E |  |  |
|  | Mean ct | 16.78 | 16.61 | 17.11 |  |  |
|  | Std. Dev | 0.180 | 0.301 | 0.056 |  |  |
|  | SEM | 0.127 | 0.213 | 0.032 |  |  |

Key: ANOVA- Analysis of Variance, SS- the sum of the square, DF- degree of freedom, SEM- standard error of the mean, EtOH-E (ethanol extract), and MeOH-E (methanol extract).

**Figure 6.**
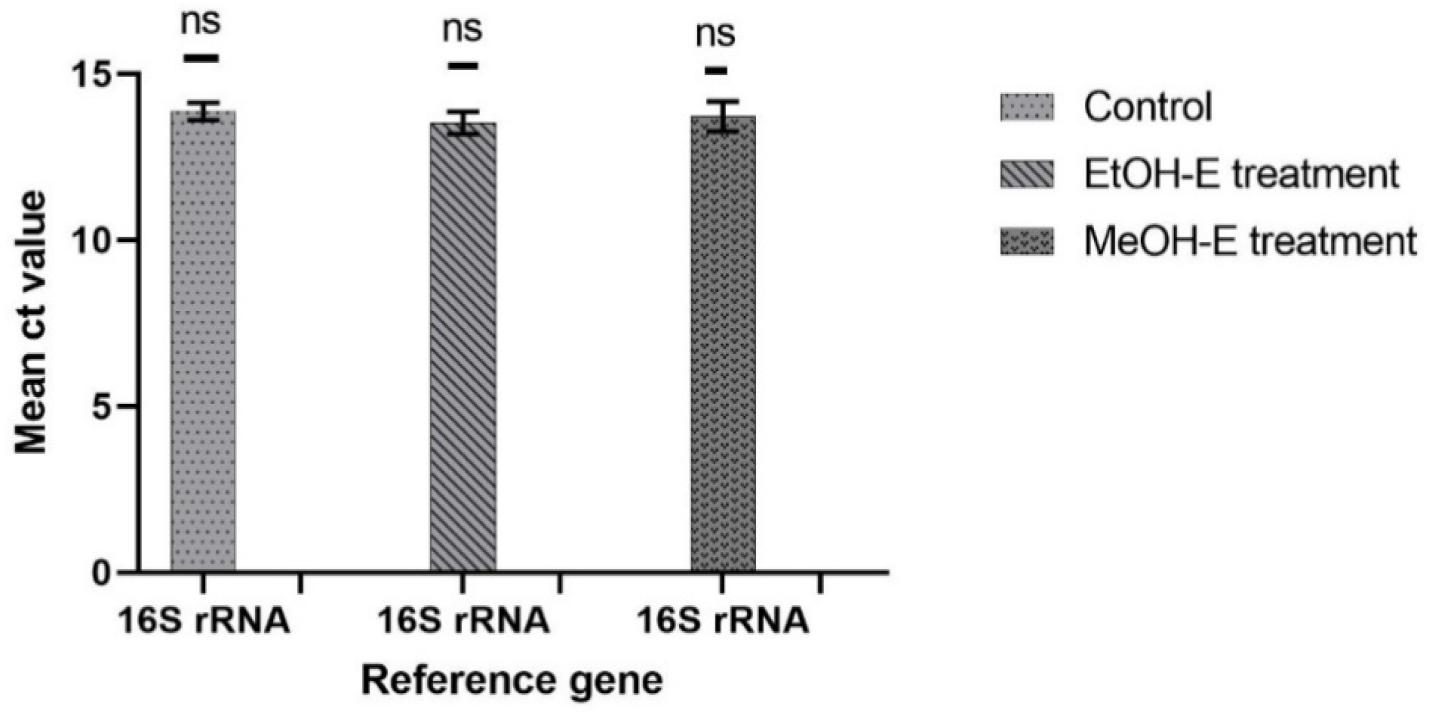
Statistical variation of the mean CT values of 16S rRNA in control, EtOH-E treatment, and MeOH-E treatment from RT-qPCR. *Key: ns-not significant, error bars-mean of triplicates ±SD (n=3), EtOH-E (ethanol extract), and MeOH-E (methanol extract)*.

**Figure 7.**
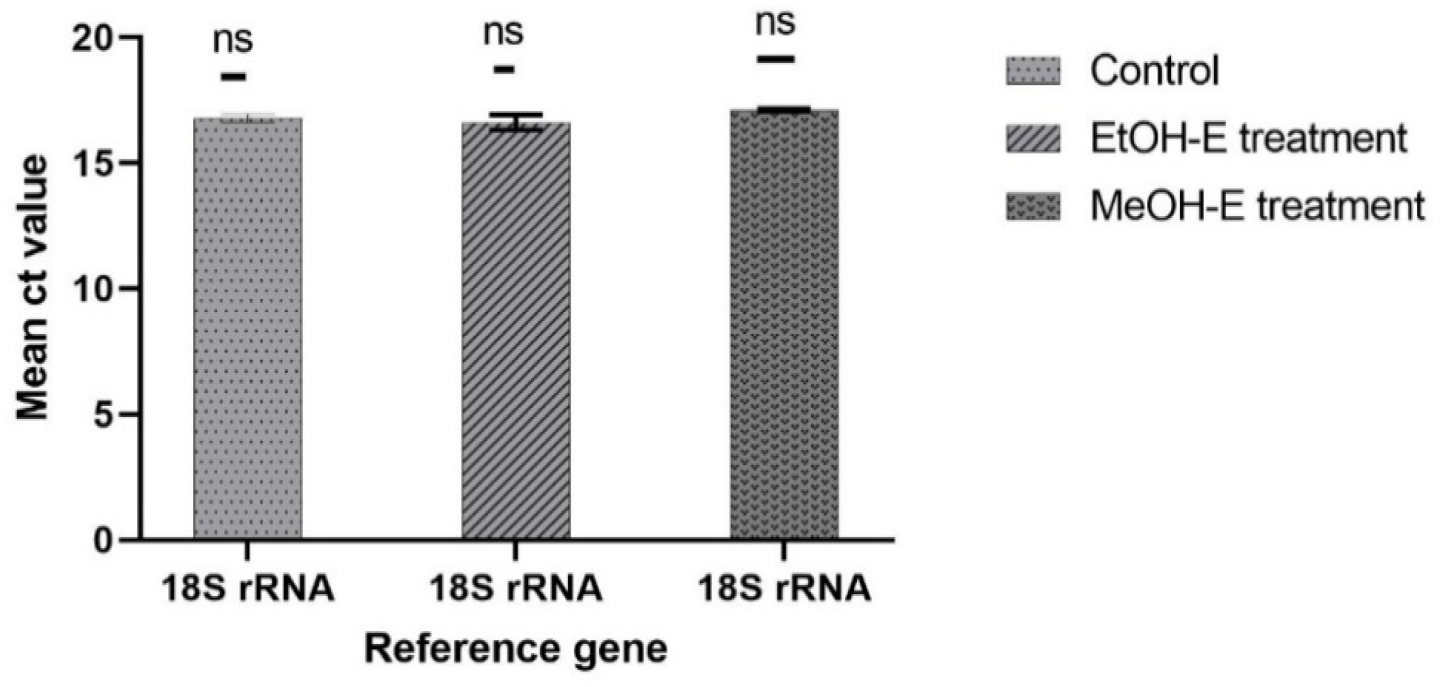
Statistical variation of the mean CT values of 18S rRNA in control, EtOH-E treatment, and MeOH-E treatment from RT-qPCR. *Key: ns-not significant, error bars-mean of triplicates ±SD (n=3), EtOH-E (ethanol extract), and MeOH-E (methanol extract)*.

### Methicillin Resistant *Staphylococcus Aureus* Virulence Gene Expression Analysis

The effects of *P. leubnitziae* ethanol and methanol extracts on the expression of *hly, sea*, and *icaA* genes were evaluated by real-time quantitative polymerase chain reaction (RT-qPCR), and the results are presented (**Table 6)**. The expression of target genes was normalized to 16S rRNA. The expression of *hly, sea*, and *icaA* of both ethanol and methanol treated samples was compared relative to the control. The fold change was analyzed using the 2^-ΔΔCT method. The fold change in the genes treated with ethanol extract (18.25 mg/ml) ranged between 0.94 and 26.66, while it ranged between 2.06 and 8.35 in the genes treated with methanol extract (18.25 mg/ml) (**Figure 8**). The differential expression analysis revealed no significant difference in *hly* between the ethanol and methanol extract treatments compared to the control (*p*-0.8671), (*p*-0.0574), respectively. Remarkably, there was a substantial difference in the *sea* (*p*-0.0007) and *icaA* (*p*-0.0001) genes treated with ethanol extract versus methanol extract (**Table 7**). The *hly* gene was less expressed (0.94-fold change) compared to control when subjected to ethanol extract treatment and expressed more than the control (which is normalized to a fold change of 1) when subjected to methanol extract treatment. Ethanol extract treatment highly induced the expression of *sea* gene (26.66-fold change) compared to methanol extract treatment (6.17-fold change). Genes were shown to be downregulated and upregulated. Genes that were above the 0 or + sign were upregulated, whereas those that were below the 0 or – sign were downregulated (**Figure 10**).

**Table 6.** RT-qPCR and 2^-ΔΔCT results of MRSA virulence genes.

| Gene | Sample | Mean CT | Std.deviation | Standard error | $\Delta CT$ | $\Delta\Delta CT$ | $2^{-\Delta\Delta CT}$ | Log2 |
| --- | --- | --- | --- | --- | --- | --- | --- | --- |
| *hly* | Control | 25.55 | 0.566 | 0.40 | 11.68 |  |  |  |
|  | EtOH-E treatment | 25.31 | 0.361 | 0.26 | 11.77 | 0.09 | 0.94 | -0.09 |
|  | MeOH-E treatment | 24.37 | 0.235 | 0.14 | 10.64 | -1.04 | 2.06 | -1.04 |
| *sea* | Control | 26.11 | 0.148 | 0.11 | 12.24 |  |  |  |
|  | EtOH-E treatment | 21.04 | 0.092 | 0.07 | 7.50 | -4.74 | 26.66 | 4.74 |
|  | MeOH-E treatment | 23.34 | 0.301 | 0.17 | 9.61 | -2.63 | 6.17 | 2.63 |
| *icaA* | Control | 24.36 | 0.057 | 0.04 | 10.49 |  |  |  |
|  | EtOH-E treatment | 23.18 | 0.071 | 0.05 | 9.65 | -0.85 | 1.8 | 0.85 |
|  | MeOH-E treatment | 21.16 | 0.090 | 0.09 | 7.44 | -3.05 | 8.3 | 3.05 |

Key: $\Delta$ - delta, CT- cycle threshold. EtOH-E (ethanol extract), MeOH-E (methanol extract).

**Table 7.**
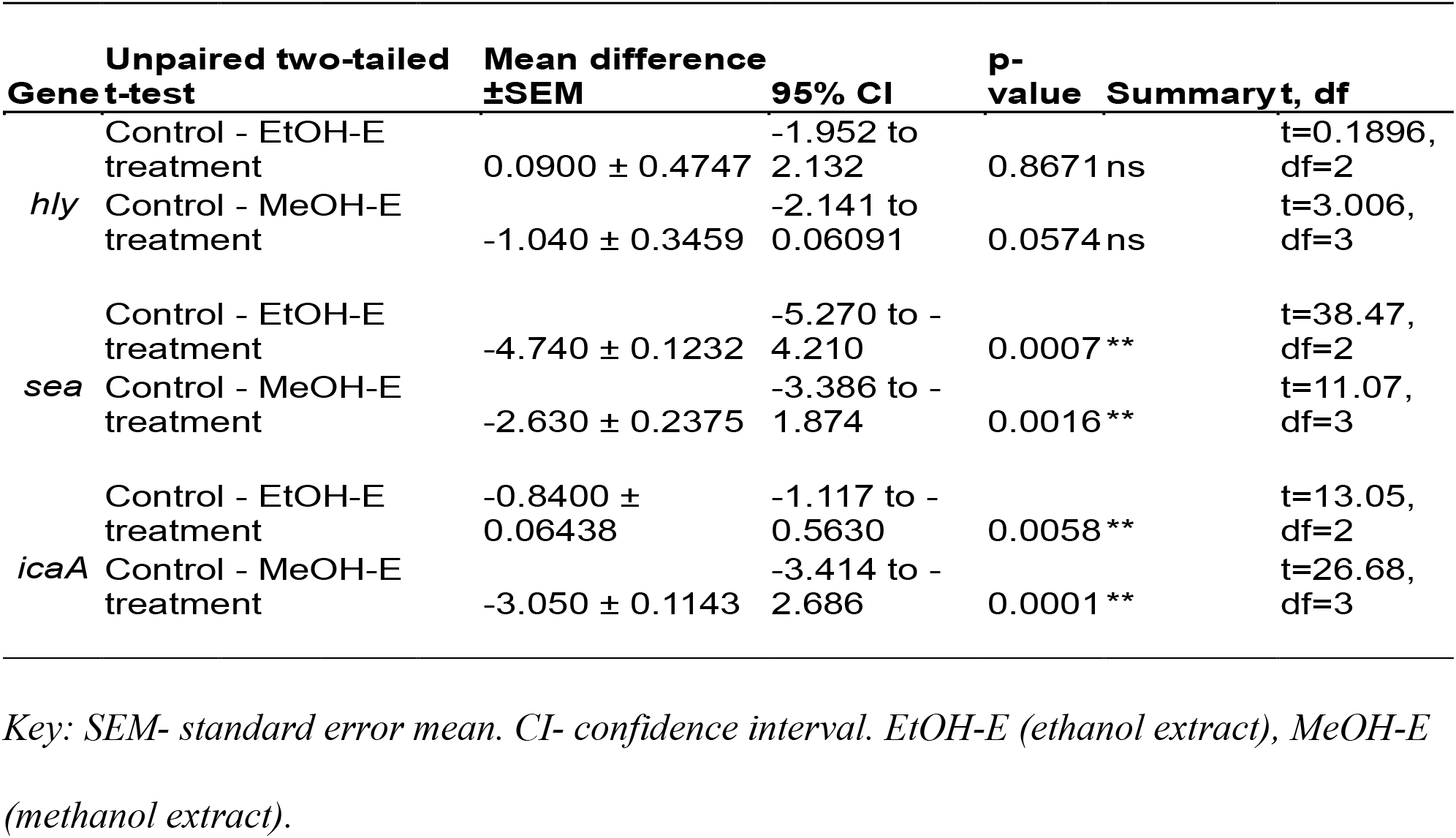
Results of unpaired two-tailed t-test on MRSA virulence genes.

**Figure 8.**
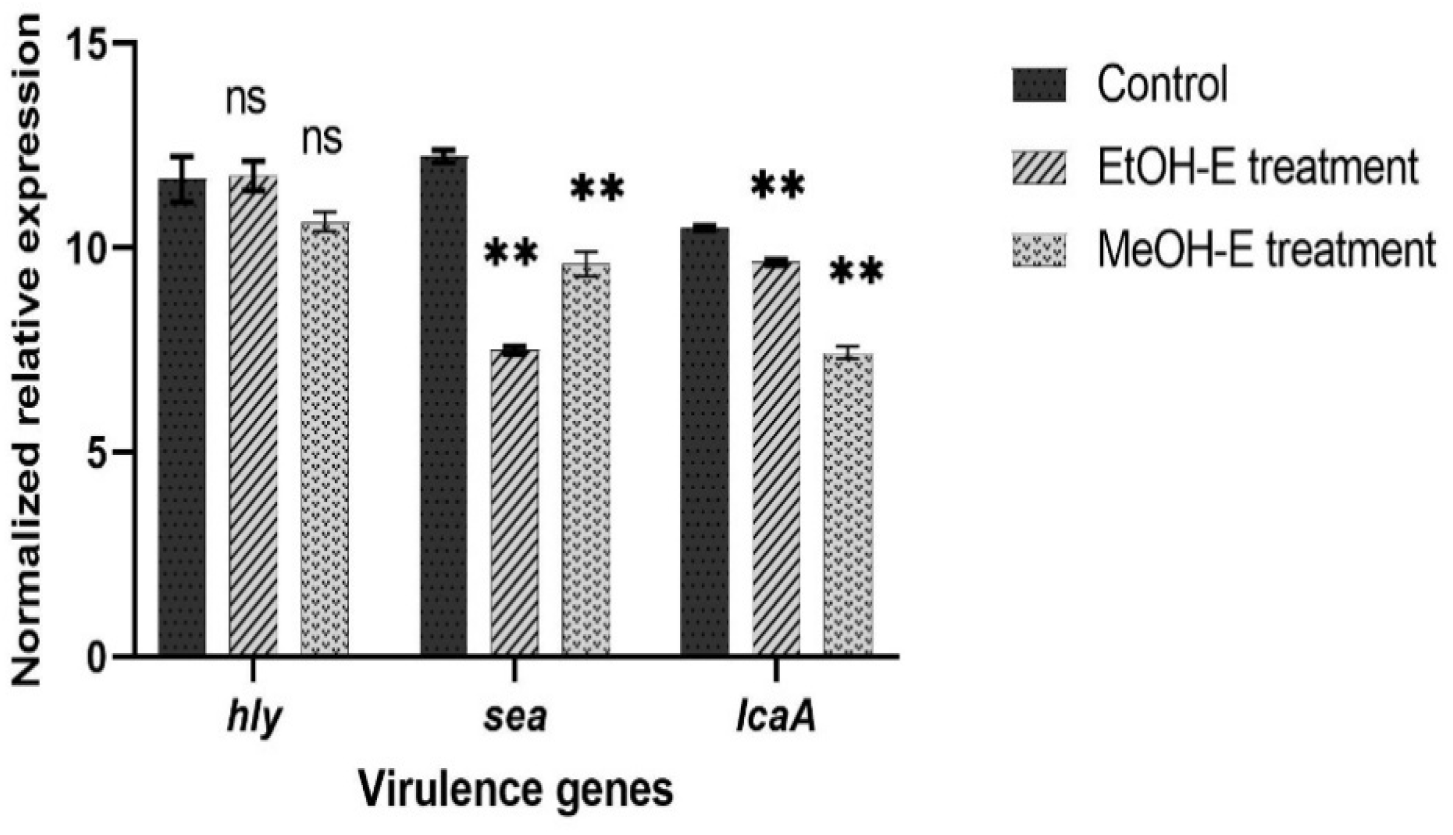
Statistical variation of ethanol and methanol extracts on the expression of MRSA virulence genes. *Key: Data are an average of ΔCT values. Bar charts represent the relative expression of hly, sea, and icaA genes treated with ethanol extract (18*.*25 mg/ml) and methanol extract (18*.*25 mg/ml) normalized with reference gene 16S rRNA and compared to the untreated (control). Error bars represent the standard deviation of triplicates. ns-not significant (p= >0*.*05), ** -significant (p= <0*.*05). Significance was compared to the control*.

**Figure 9.**
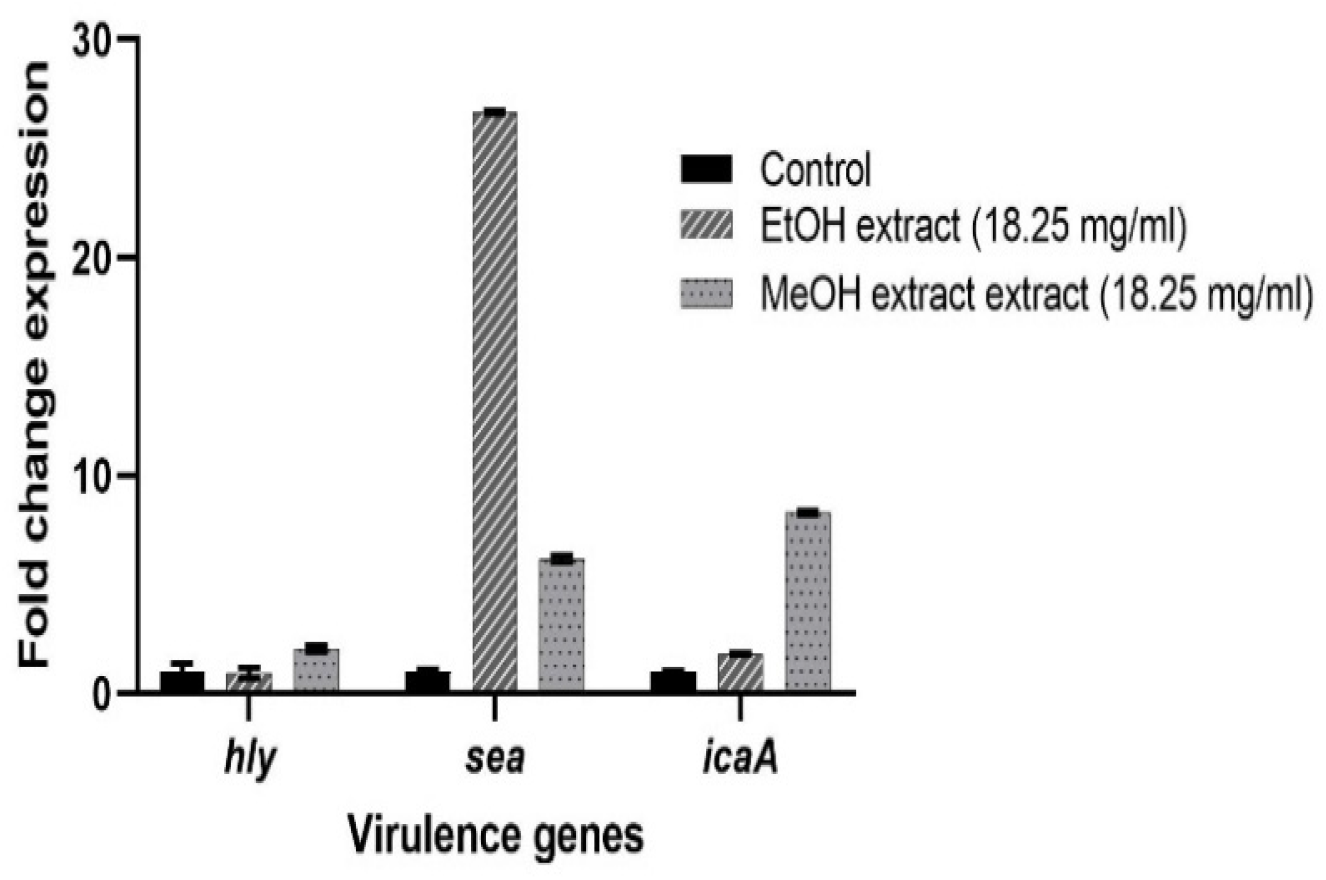
Gene expression level of MRSA virulence genes. *Key: Bar chart represents the fold change of hly, sea, and icaA genes treated with sub-MIC of ethanol extract (18*.*25 mg/mL) and methanol extract (18*.*25 mg/mL), normalized with the reference gene 16S rRNA and compared to the untreated calibrator. The error bars indicate the standard error mean*.

**Figure 10.**
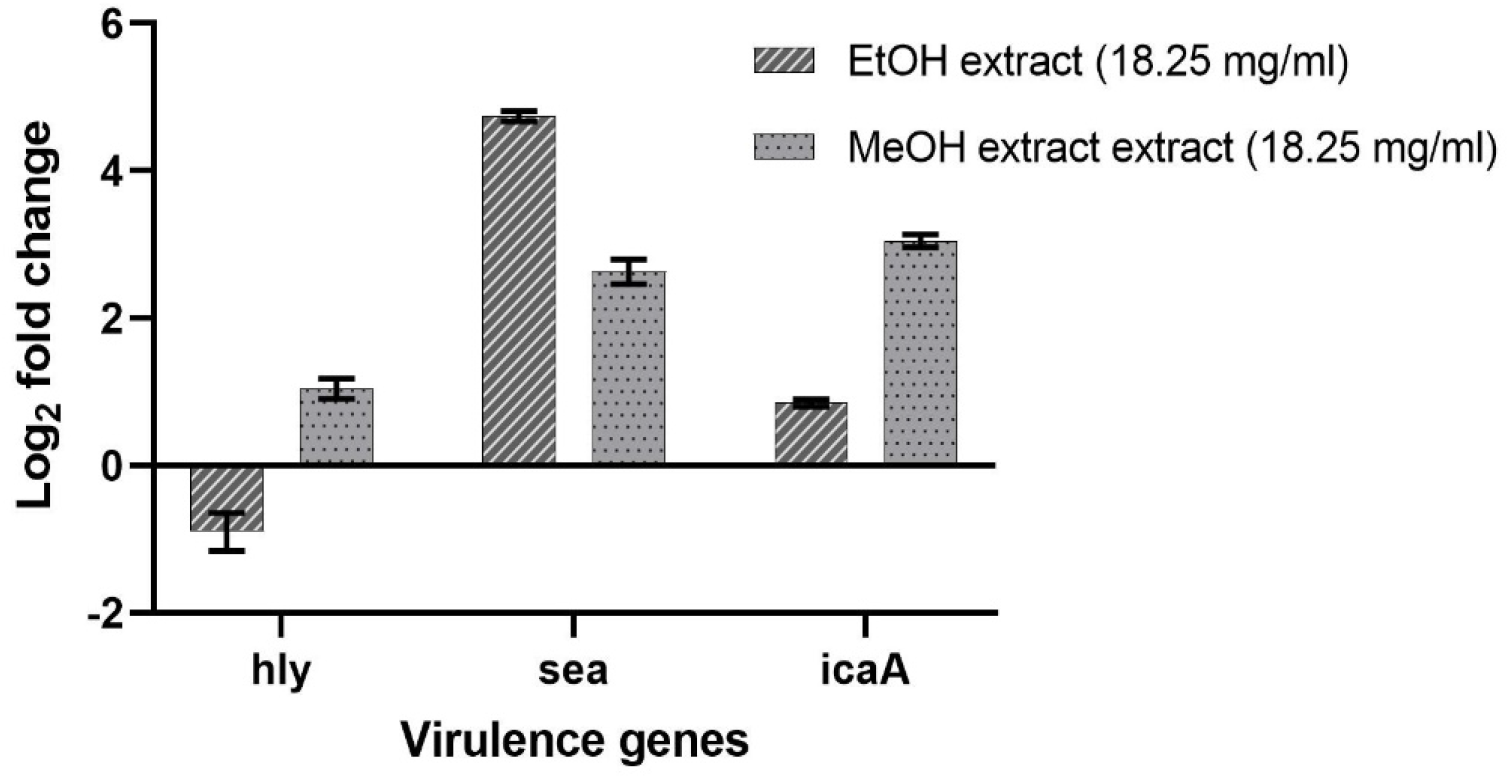
Effect of *P. leubnitziae* ethanol and methanol extracts on the expression of virulence genes of MRSA. *Key: Bar charts represent the fold change log2 transformed to show the downregulation or upregulation of hly, sea, and icaA genes treated with sub-MIC of ethanol extract (18*.*25 mg/mL) and methanol extract (18*.*25 mg/mL). The error bars indicate the standard error mean*.

### *Candida albicans* ATCC 90029 Virulence Gene Expression Analysis

The effects of *P. leubnitziae* ethanol and methanol extracts on the expression of *als1, pbl1*, and *sap1* genes were evaluated by real-time quantitative polymerase chain reaction (RT-qPCR), and the results are presented (**Table 8)**. The expression of target genes was normalized to 18S rRNA. The expression of *als1, pbl1*, and *sap1* of both ethanol and methanol treated samples was compared relative to the control. The fold change was analyzed using the 2^-ΔΔCT method. The fold change ranged between 0.00001 – 0.004 in the genes treated with ethanol extract (2.344 mg/ml) while ranged between 0.002 – 0.003 in the genes treated with methanol extract (9.375 mg/ml) (**Figure 11**). The differential expression analysis showed a significant difference of all genes of the ethanol and methanol extracts treatment to that of the control. The ethanol extract treatment of *als1, pbl1*, and *sap1* exhibited a *p*-value of 0.0001, 0.0001, and 0.0002, respectively. For methanol extract treatment, the genes had lower *p*-values of <0.0001 (**Table 9)**. All genes when subjected to ethanol and methanol extracts treatment were less expressed, denoting a zero-fold change compared to the control (normalized to 1-fold change) (**Figure 12)**. Notably, the genes subjected to both treatments showed a negative log2 which validates a downregulation (**Figure 13**).

**Table 8.**
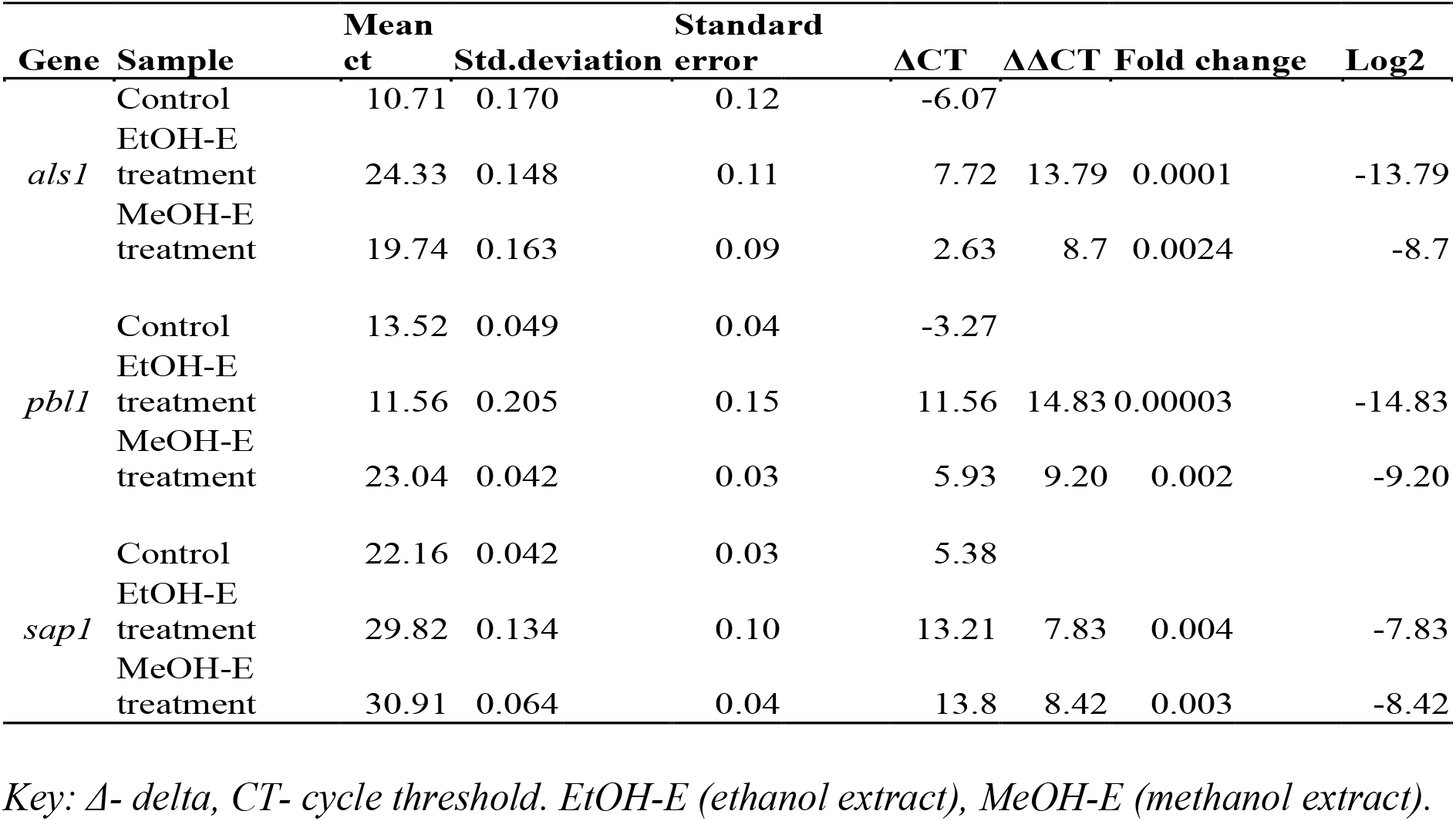
RT-qPCR and 2^-ΔΔCT results of *Candida albicans* ATCC 90029 virulence genes.

| Gene | Sample | Mean<br>ct | Std.deviation | Standard<br>error | $\Delta CT$ | $\Delta\Delta CT$ | Fold change | Log2 |
| --- | --- | --- | --- | --- | --- | --- | --- | --- |
| *als1* | Control | 10.71 | 0.170 | 0.12 | -6.07 |  |  |  |
|  | EtOH-E<br>treatment | 24.33 | 0.148 | 0.11 | 7.72 | 13.79 | 0.0001 | -13.79 |
|  | MeOH-E<br>treatment | 19.74 | 0.163 | 0.09 | 2.63 | 8.7 | 0.0024 | -8.7 |
| *pbl1* | Control | 13.52 | 0.049 | 0.04 | -3.27 |  |  |  |
|  | EtOH-E<br>treatment | 11.56 | 0.205 | 0.15 | 11.56 | 14.83 | 0.00003 | -14.83 |
|  | MeOH-E<br>treatment | 23.04 | 0.042 | 0.03 | 5.93 | 9.20 | 0.002 | -9.20 |
| *sap1* | Control | 22.16 | 0.042 | 0.03 | 5.38 |  |  |  |
|  | EtOH-E<br>treatment | 29.82 | 0.134 | 0.10 | 13.21 | 7.83 | 0.004 | -7.83 |
|  | MeOH-E<br>treatment | 30.91 | 0.064 | 0.04 | 13.8 | 8.42 | 0.003 | -8.42 |

Key: $\Delta$ - delta, CT- cycle threshold. EtOH-E (ethanol extract), MeOH-E (methanol extract).

**Table 9.**
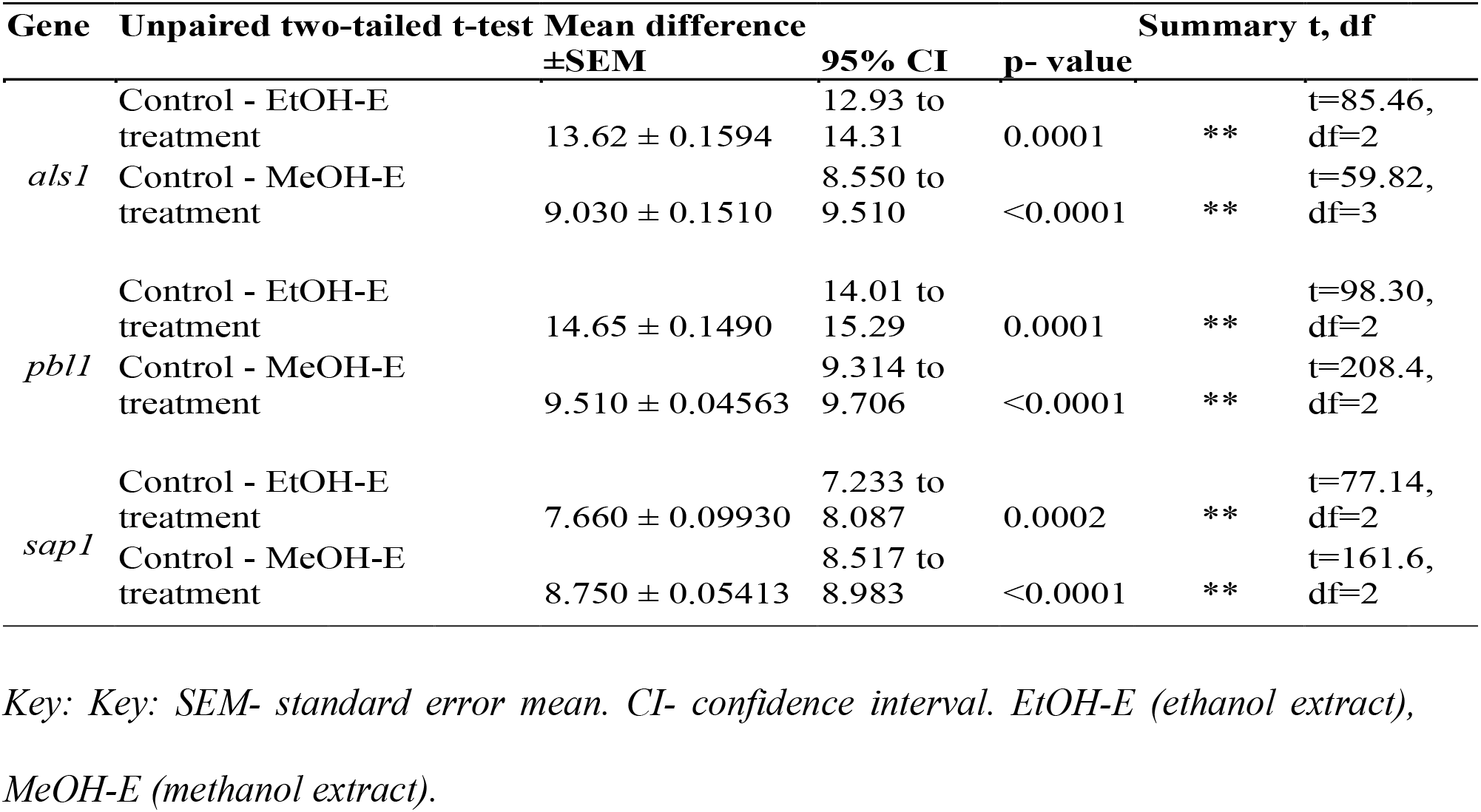
Results of unpaired two-tailed t-test on *Candida albicans* ATCC 90029 virulence genes.

**Figure 11.**
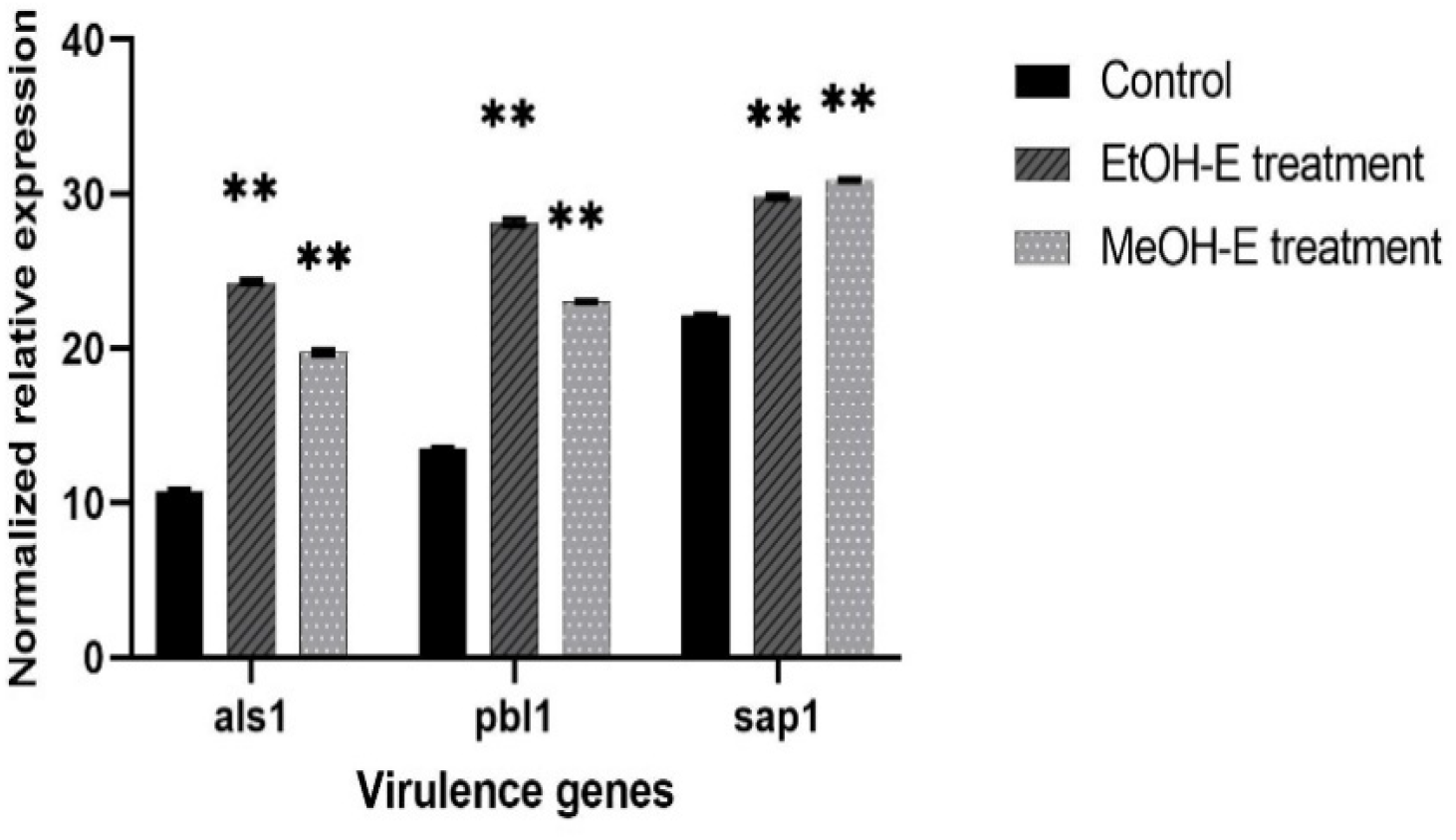
Statistical variation of *P. leubnitziae* ethanol and methanol extracts on the expression of *C. albicans* ATCC 90029 virulence genes. *Key: Data are an average of ΔCT values. Bar charts represent the relative expression of als1, pbl1, and sap1 genes treated with ethanol extract (2*.*344 mg/ml) and methanol extract (9*.*375 mg/ml) normalized with reference gene 18S rRNA and compared to the untreated (control). Error bars represent the standard deviation of triplicates. ** -significant (p= <0*.*05). Significance was compared to the control*.

**Figure 12.**
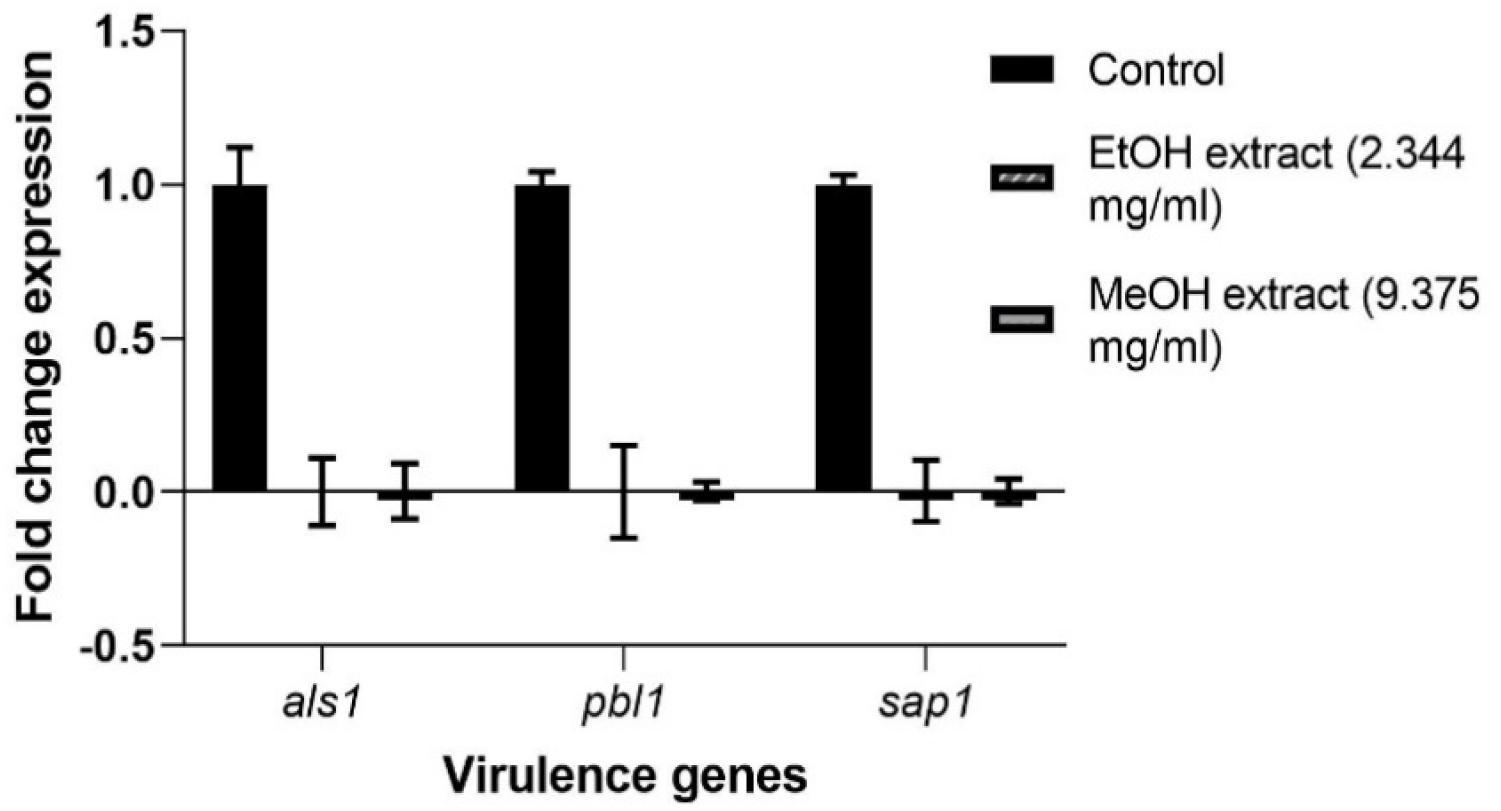
Gene expression level of *C. albicans* ATCC 90029 virulence genes *Key: Bar chart represents the fold change of als1, pbl1, and sap1 genes treated with sub-MIC of ethanol extract (2*.*344 mg/mL) and methanol extract (9*.*375 mg/mL), normalized with the reference gene 18S rRNA and compared to the untreated calibrator. The error bars indicate the standard error mean*.

**Figure 13.**
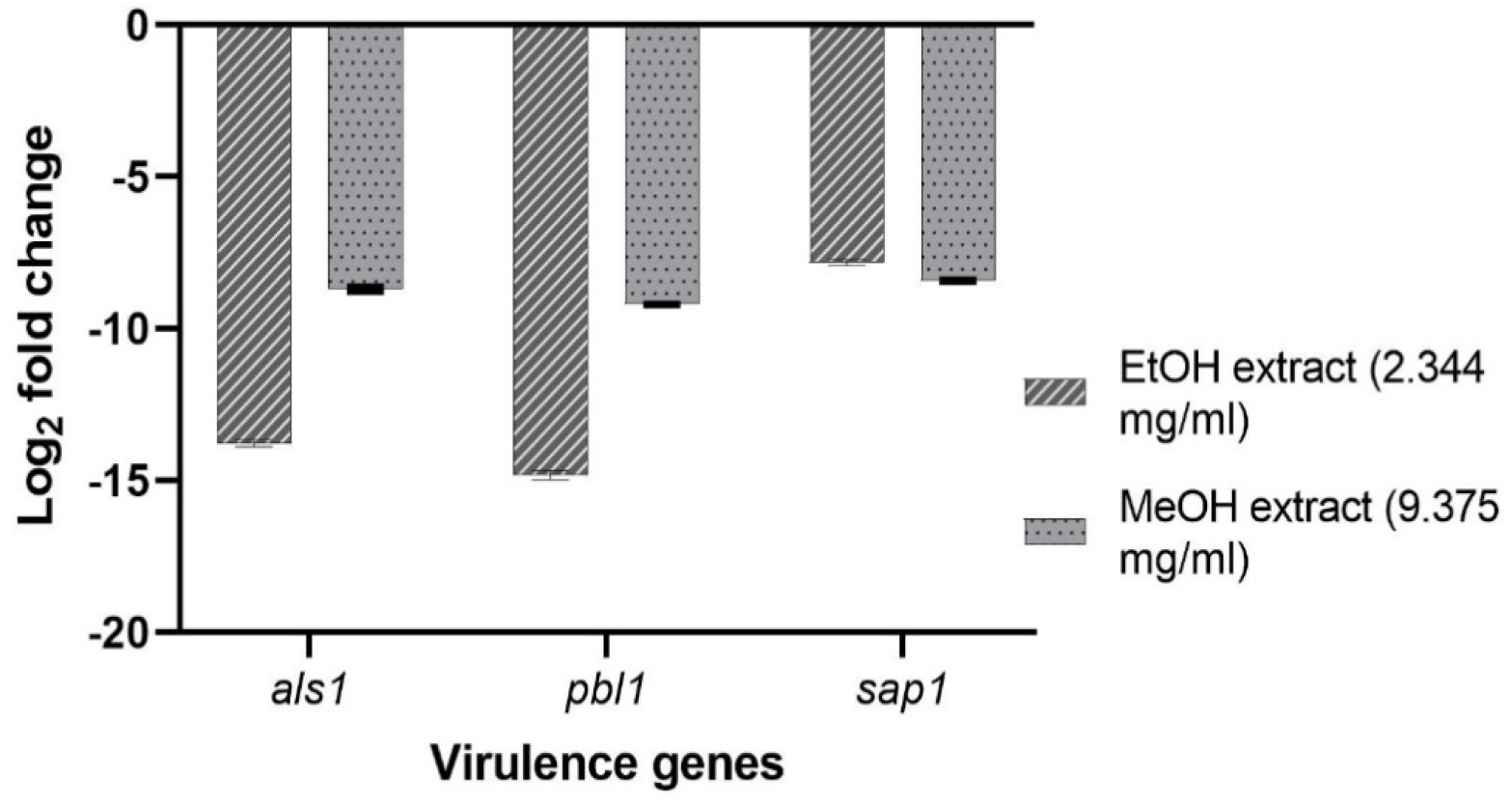
Effect of *P. leubnitziae* ethanol and methanol extracts on the expression of virulence genes of *C. albicans* ATCC 90029. *Key: Bar charts represent the fold change log2 transformed to show downregulation or upregulation of als1, pbl1, and sap1 genes treated with sub-MIC of ethanol extract (2*.*344 mg/mL) and methanol extract (9*.*375 mg/mL). The error bars indicate the standard error mean*.

## DISCUSSION

The leaf extract of *Pechuel leubnitziae* was investigated in this study. The rising number of cases associated with antimicrobial resistance is currently a major public health concern [1]. Antimicrobial resistance has been attributed to the burden of managing recurrent infections, prolonged hospitalization, and an increased morbidity and mortality rate [1]. The antimicrobial activity of ethanol and methanol extracts against clinical isolates associated with antimicrobial resistance was investigated. The current study’s findings revealed that ethanol and methanol extracts have antimicrobial activity, inhibiting 40% of the test organisms. The extracts were more effective against the fungus, exhibiting antimicrobial activity at all concentrations (200-600 mg/ml). The results concur with the antimicrobial activity of *P. leubnitziae* methanol root extract observed by [15] and the antibacterial activity reported by [16]. The findings of the study contrast to that of [15], which showed no antimicrobial activity against *S. aureus* and *Candida albicans* adopting the disc diffusion method. The variations in the results of *P. leubnitziae* leaf extract could be attributed to the concentrations of 2.5 mg/ml, 5 mg/ml, and 10 mg/ml used by [15], as opposed to the current study, which used concentrations of 200 mg/ml, 400 mg/ml, and 600 mg/ml. It could also have been due to the antimicrobial susceptibility assay used in each study. The current study has used the agar well diffusion method, which allows for high volume disperse of the test solution in the well versus the disc diffusion method used by [15], which only enables very low absorbance of the test solution by the disc. The antimicrobial effectiveness of plant crude extract is greatly proportional to the dose and strains; this could explain why the current study’s findings differ from those published in a prior study, which used lower concentrations [17].

The antimicrobial activity of the ethanol and methanol extracts was verified using 5% DMSO alone, indicating that the action was exhibited by the extracts rather than the solvent employed to reconstitute the extracts. Antibacterial activity was exclusively observed in gram-positive bacteria, and without any activity observed in gram-negative bacteria. Gram-positive bacteria have a thicker cell wall that is rich in peptidoglycan and teichoic acid. Gram-negative bacteria, on the other hand, have an outer membrane and a periplasmic region rich in peptidoglycan, as well as an internal lipopolysaccharide layer. As a result, whereas compounds from *P. leubnitziae* leaf extract may be able to diffuse well through the outer wall of Gram-positive bacteria, they may not be able to diffuse effectively through all of the narrow channels of the Gram-negative outer membrane [18].

According to the [13] recommendation, at least half of the antibiotics used failed to achieve the required inhibitory value for reactivity, primarily against gram-positive test organisms (Figure 1). This could underlie the selective activity of *Pechuel-Loeschea leubnitziae* leaf extract. The MIC of the ethanolic extract for *Candida* was 4.688 mg/ml and 37.5 mg/ml for MRSA, while the MIC of the methanolic extract was 18.75 mg/ml for *Candida* and 37.5 mg/ml for MRSA. A high MIC value indicates poor activity, whereas a low MIC value indicates superior activity. In the current investigation, the ethanol extract showed a lower MIC against *Candida* than the methanol extract, implying that it is more potent. At 37.5 mg/ml, the bacteriolytic effect of the ethanol and methanol extracts was observed. Additionally, the fungistatic and fungicidal activities of ethanol and methanol were found to be 18.75 mg/ml and 37.5 mg/ml, respectively. At a low concentration, the ethanol extract exhibited a lethal effect, revealing preliminary evidence on its consistency as the most effective potential source of antimicrobial compounds in the treatment of microbial disorders. This observation of MIC values lines with the observations of [16] and [19]. Another study found that *Streptomyces* spp. SRF1 culture filtrate extract had a minimum bactericidal concentration of (0.390 mg/ml) against MRSA [20]. RT-qPCR was used in this project to assess the influence of *P. leubnitziae* ethanol and methanol extracts on the expression of *hly, sea*, and *icaA*. Emerging strategies rely on the therapeutic potential of natural compounds on the pathogenesis mechanisms of microbes. Endogenous genes were used as positive controls to validate and normalize the RT-qPCR data. The difference in the mean CT values of endogenous expression between treated and untreated samples should not be statistically significant. According to the current study’s findings, exposing MRSA to a sub-MIC concentration (18.25 mg/ml) of ethanol extract had an inhibitive effect on the expression of *hly*, resulting in a downregulation. The findings are consistent with the findings of [21], who reported a reduction in *hly* and *flu* gene expression when MRSA is exposed to D-SNPs and silver nitrate. Another study, carried out by [22], discovered that *Rhus javanica* extract completely suppressed *sea* expression.

The alpha-hemolysin gene is a crucial gene from hemolysin virulence factor of MRSA. The downregulation in the expression of the *hly* gene when treated with sub-MIC of the ethanol extract of *P. leubnitziae* leaves could be indicative that it influences the bacterial defense mechanisms by lessening virulence and invasiveness [23, 21]. These findings rule out the possibility of the ethanol extract having a crucial anti-virulence effect on the expression of the *hly* gene. The formation of biofilm and release of toxins are reported to be influenced by various conditions that are likely toxic, including an excessive amount of urea, osmolality, detergents, oxidative stress, and sub-MICs. The upregulation of *sea* and *icaA* genes, when exposed to both sub-MIC of ethanol and methanol extracts and *hly* gene when exposed to methanol extract, could be the pathogen’s survival mechanism in response to adverse [24, 21]. Thus, showing no attenuation effect of the extracts on the upregulated genes.

To allow the quantification of the expression of *C. albicans* ATCC 90029, *als1, pbl1*, and *sap1* virulence genes treated with sub-MIC of *P. leubnitziae* ethanol extract (2.344 mg/ml), methanol extract (9.375 mg/ml), and untreated, real-time qPCR was used. The virulence of *Candida albicans* is attributed to several factors, including adhesion, invasion, and protease production. These virulence factors play an essential role in the pathogenesis of *C. albicans* [25]. This study reported a significant downregulation in the expression of *als1, pbl1*, and *sap1* genes when exposed to ethanol and methanol extracts. These findings are similar to the data reported by [25], who reported a suppression in *sap1* and *sap10* when treated with *Pluchea dioscoridis*. Another study done by [26], reported a significant downregulation on the expression level of *als1, als2, hwp1, SapT1, SapT2*, and *plb1* genes when treated with Morin. The differential expression that showed significant downregulation of *C. albicans* ATCC 90029 virulence genes could be due to the presence of phenolics compounds reported from quantitative analysis of *P. leubnitziae* leaf extract of ethanol and methanol [27]. Phenolics compounds are believed to influence the apoptotic pathway against *C. albicans* [28].

## CONCLUSION

The antimicrobial activity of *P. leubnitziae* leaf extract against tested organisms affirms its traditional application in the treatment of microbial infections, per the findings of this study. Ethanol extract had greater fungicidal activity against *Candida albicans*, suggesting that it could be an effective and prospective source of antifungal agents in herbal formulations. When *C. albicans* ATCC 90029 was treated with sub-MIC (2.344 and 9.375 mg/ml) of ethanol and methanol extracts, gene expression analysis revealed a substantial downregulation of the *als1, pbl1*, and *sap1* genes. In conclusion, ethanol extract suppressed the expression of the *hly* gene in MRSA, resulting in a downregulation. Furthermore, the ethanol and methanol extracts had no influence on the *sea* and *icaA* genes in MRSA. Overall, the current study discloses the potential of ethanol and methanol extracts of *P. leubnitziae* leaf extract as an effective potential source of antimicrobial agents in the treatment of MRSA and *C. albicans* related infections.

## RECOMMENDATION

More research is needed to assess the antimicrobial activity of various solvents. It is possible to develop and evaluate antifungal gels and creams using ethanol extract. additionally, *in-vivo* therapeutic efficacy investigations, as well as the identification and purification of pure bioactive compounds from *P. leubnitziae* leaves, will be crucial for using this plant leaf extract in drug development and discovery. Molecular docking studies of bioactive compounds identified in P. leubnitziae leaves could be used to confirm the binding affinity of the compound to the protein molecules of MRSA and *Candida albicans* generated from various major virulence factors.

## ACKNOWLEDGMENT

We would like to applaud the Pan African University of Basic Science, Technology, and Innovation for providing infrastructure, as well as Ifeoluwa Deborah Gbala for laboratory guidance all through the completion of the work. The authors are immensely thankful to the African Union for financing their research.

